# Identification of a specific biomarker of *Acinetobacter baumannii* Global Clone 1 by machine learning and PCR related to metabolic fitness of ESKAPE pathogens

**DOI:** 10.1101/2021.10.18.464923

**Authors:** Verónica Elizabeth Álvarez, María Paula Quiroga, Daniela Centrón

**Affiliations:** Laboratorio de Investigaciones en Mecanismos de Resistencia a Antibióticos (LIMRA), Instituto de Investigaciones en Microbiología y Parasitología Médica, Facultad de Medicina, Universidad de Buenos Aires - Consejo Nacional de Investigaciones Científicas y Tecnológicas (IMPaM, UBA-CONICET), Buenos Aires, Argentina; Nodo de Bioinformática. Instituto de Investigaciones en Microbiología y Parasitología Médica, Facultad de Medicina, Universidad de Buenos Aires - Consejo Nacional de Investigaciones Científicas y Técnicas (IMPaM, UBA-CONICET), Ciudad Autónoma de Buenos Aires, Argentina

## Abstract

Since the emergence of high-risk clones worldwide, constant investigations have been undertaken to comprehend the molecular basis that led to their prevalent dissemination in nosocomial settings over time. So far, the complex and multifactorial genetic traits of this type of epidemic clones have only allowed the identification of biomarkers with low specificity. A machine learning algorithm was able to recognize unequivocally a biomarker for the early and accurate detection of *Acinetobacter baumannii* Global Clone 1 (GC1), one of the most disseminated high-risk clones. Support Vector Machine identified the U1 sequence with 367 nucleotides length that matched a fragment of the *moaCB* gene, which encodes the molybdenum cofactor biosynthesis C and B proteins. U1 differentiates specifically between *A. baumannii* GC1 and non-GC1 strains, becoming a suitable biomarker capable of being translated into clinical settings as a molecular typing method for early diagnosis based on PCR as shown here. Since the metabolic pathways of Mo enzymes have been recognized as putative therapeutic targets for ESKAPE pathogens, our findings highlighted that machine learning can be also useful in intricate knowledge gaps of high-risk clones and implies noteworthy support to the literature to identify challenging nosocomial biomarkers for other multidrug-resistant high-risk clones.

**IMPORTANCE:** *A. baumannii* GC1 is an important high-risk clone that rapidly develops extreme drug resistance in the nosocomial niche. Furthermore, several strains were identified worldwide in environmental samples exacerbating the risk of human interactions. Early diagnosis is mandatory to limit its dissemination and to outline appropriate antibiotic stewardship schedules. A region of 367 bp length (U1) within the *moaCB* gene not subjected to Lateral Genetic Transfer or to antibiotic pressures was successfully found by Support Vector Machine algorithm that predicts *A. baumannii* GC1 strains. PCR assays have confirmed that U1 specifically identifies *A. baumannii* GC1 strains. At the same time, research on the group of Mo enzymes proposed this metabolic pathway related to superbuǵs metabolism as a potential future drug target site for ESKAPE pathogens due to its central role in bacterial fitness during infection. These findings confirmed the importance of machine learning applied to the burden of the rise of antibiotic resistance.

## INTRODUCTION

*Acinetobacter baumannii* is an opportunistic and nosocomial Gram-negative pathogen that causes a wide range of nosocomial infections. It is included in the group of ESKAPE pathogens (1, 2). Nosocomial infections by this species have increased in recent years, which added to its ability to acquire and spread antibiotic resistance genes to all families of antibiotics, are considered a serious global threat worldwide (3, 4). The majority of *A. baumannii* isolates that are broadly resistant to antibiotics belong to two pandemic clones, known as global clones 1 (GC1) and 2 (GC2) (3, 5, 6). Recent epidemiological studies of carbapenem-resistant *A. baumannii* (CRAB) isolates, revealed that *A. baumannii* GC1 is the prevalent CRAB clone in several countries (7–10). In addition, *A. baumannii* GC1 strains have been found worldwide in environmental samples from water, soil, and animals (11, http://www.acinetobacterbaumannii.no/).

Over time, molecular methods with different degrees of resolution have been used to type *A. baumannii* strains such as amplified fragment length polymorphism analysis, ribotyping, macrorestriction analysis by pulsed-field gel electrophoresis, multiplex polymerase chain reactions (PCR), multilocus sequence typing (MLST) and more recently whole genome sequencing (WGS) (12–18). A typing scheme based on two multiplex PCR targeting three genes (*ompA*, *csuE*, and *bla*_OXA-51-like_) has been used for the assignment of *A. baumannii* isolates into two major PCR-based groups corresponding to *A. baumannii* GC1 and GC2 (19). Also, since correlation has been detected between particular *bla*_OXA-51- like_ alleles and some epidemic lineages, sequence analysis of the *bla*_OXA-51-like_ gene has been proposed as a useful typing method for *A. baumannii* isolates (19–21). In agreement to this, a study conducted on sixty *A. baumannii* isolates collected worldwide demonstrated that isolates belonging to *A. baumannii* GC1 encoded enzymes from the OXA-69 cluster which included OXA-69, OXA-92, OXA-107, OXA-110 and OXA-112 enzymes (20). However, not all the isolates encoding the OXA-69 cluster belonged to *A. baumannii* GC1 such as *A. baumannii* A92 strain that belongs to GC2 (20). An additional typing method of *A. baumannii* GC1 strains, consisted in the detection of 108 bp deletion in the 5′-conserved segment (5′-CS) of the class 1 integron located in AbaR3 genomic island (22). Nevertheless, since AbaR3 is not present in all *A. baumannii* GC1 strains, this approach serves as a marker for some diverged lineages within *A. baumannii* GC1 clone (22). All the previously described methods based on PCR include target genes that are subjected to Lateral Genetic Transfer and/or antibiotic pressure, which represents a limitation for the specificity of the technique.

With the increasing throughput and decreasing cost of DNA sequencing, large numbers of bacterial genomes have been submitted to public databases (23, 24). Genome-wide studies of DNA variation related to antibiotic resistance phenotypes have garnered high public interest, especially since several multidrug-resistant strains have emerged worldwide (25). New candidate biomarkers leading to the identification of resistant pathogens require the study and development of fast, easily applicable, and accurate tools. Furthermore, with the help of computational algorithms, such studies can be conducted at a much larger scale producing more significant results (26–29). Machine learning (ML) algorithms and statistics have been used increasingly to build models that correlate genomic variations with phenotypes that may help to predict bacterial phenotypes and genotypes (30–36). In supervised ML, each learning sample includes the outcome (class label) of interest, and it is used to build a prediction model (37). The model takes an outcome measurement (e.g., a bacterium having a resistant phenotype or genotype) and tries to learn from the available data (e.g., information on genomic mutations) to predict the outcome measure. The developed model is then applied to new and unseen data. The goal of the algorithm is to train a model that accurately foresees the correct outcome for any input (38).

Support Vector Machine (SVM) is a supervised learning algorithm formally characterized by a separating hyperplane that divides binary data to solve both classification and regression problems. SVM aims to correctly classify samples based on examples in the training dataset (39–41). On the other hand, Set Covering Machine (SCM) is a supervised learning algorithm that uses a greedy approach to produce uncharacteristically sparse rule-based models from the input data (42). Both algorithms have been applied to several biological knowledge gaps and proved to be accurate in predicting novel antibacterial agents (43), antibiotic resistance genes (32, 33, 44–46), identification of microorganisms (47–49), cancer diagnosis (50–53), among others (30, 47, 54–57).

ML can deal with large and diverse datasets to extract relevant information (58). Given the wide use of ML in biology, and the absence of accurate identifiers for early diagnosis of *A. baumannii* GC1 strains, our study aimed to assess whether ML could be applied to process thousands of genomes to identify a suitable *A. baumannii* GC1 biomarker. We also wanted to analyze whether ML could be combined with other techniques such as PCR and/or qPCR with high-resolution melting (HRM) assays to provide a molecular typing method capable of being translated into clinical settings. For these reasons, we built predictive models for typing *A. baumannii* GC1 genomes by training SVM and SCM classifiers. From these classifiers, we identified a new and specific genomic biomarker for the early detection of *A. baumannii* GC1 strains by PCR technique not subjected to selective pressure of antibiotics nor to Lateral Genetic Transfer.

## RESULTS

To identify new genomic biomarkers that uniquely identify strains belonging to *A. baumannii* GC1, we applied the SVM and SCM algorithms to the Dataset 1 (500 genomes) and the Dataset 2 (4799 genomes) (Table S1 and S2). Firstly, we applied both algorithms to the Dataset 1 which was composed of 200 *A. baumannii* GC1 and 300 *A. baumannii* non- GC1 genomes. We aimed to predict whether a particular genome in the Dataset 1 belonged to *A. baumannii* GC1 or not by using SVM and SCM algorithms. The Dataset 1 was used as input for both algorithms during the training and the testing of the models. Once we obtained accurate models that predicted putative biomarkers sequences, we used the second *A. baumannii* genome collection, the Dataset 2 (Table S2). The Dataset 2 was composed of 312 *A. baumannii* GC1 and 4487 *A. baumannii* non-GC1 genomes. By using blastn searches, we analyzed whether the predicted putative biomarker sequences were also found in the Dataset 2 genomes maintaining the same pattern found in the Dataset 1. This analysis was done to perform an external validation of the predictions made by both algorithms. The study workflow is summarized in Figure 1.

**Figure 1.**
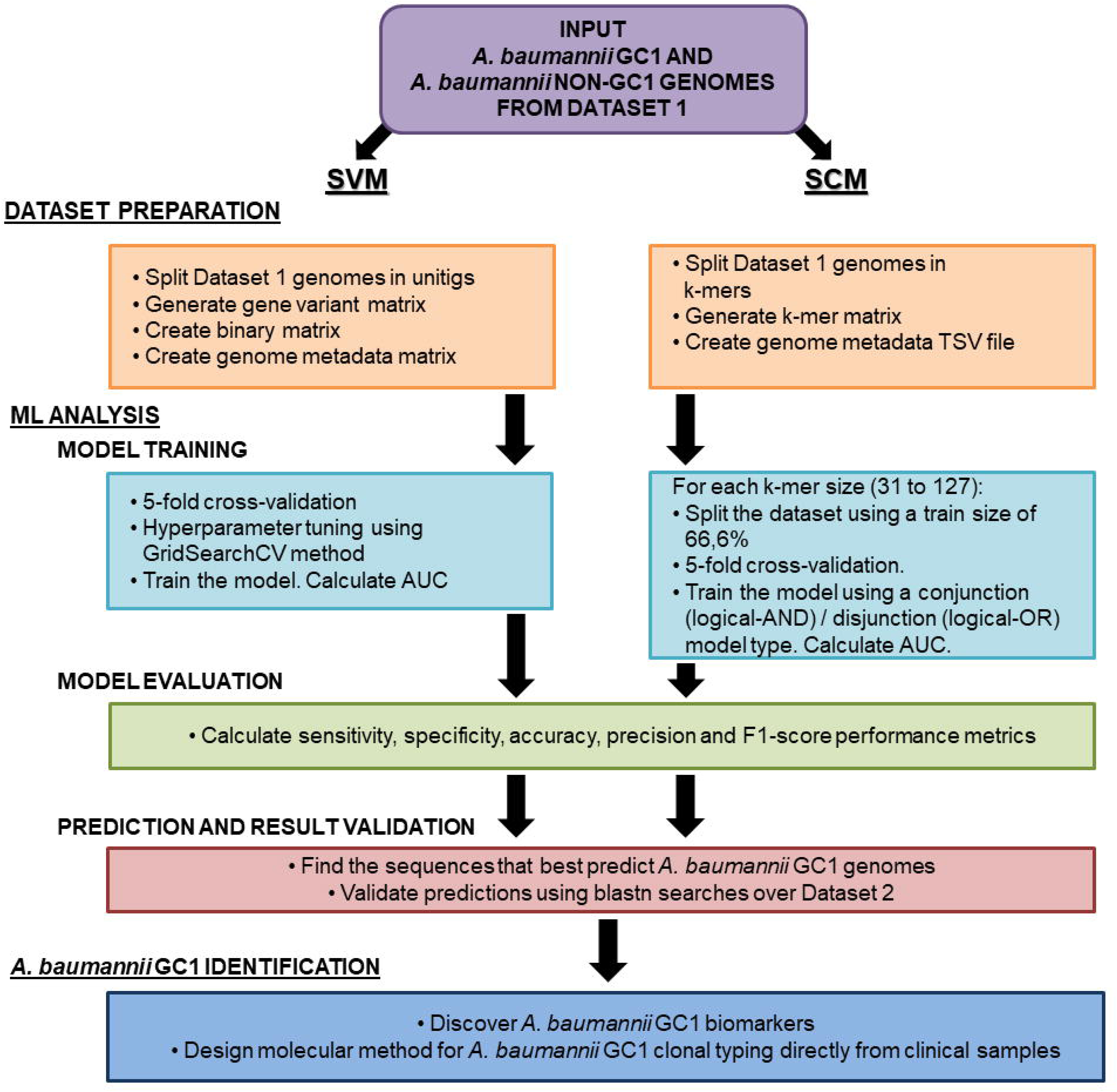
Diagram of the workflow for *A. baumannii* GC1 biomarker discovery using machine learning. The diagram indicates the steps we followed during the present work related to genomes collections used, dataset preparation, ML analysis and the design of a method for *A. baumannii* GC1 strain identification.

### Obtaining unique *A. baumannii* GC1 predictive sequences with SVM

The Pasteur scheme for Multilocus Sequence Typing (MLST) has the potential to identify isolates belonging to *A. baumannii* GC1 providing a neat demarcation of STs composing GC1 and non-*A. baumannii* clonal complexes (5, 59–62). Considering this fact, we annotated the ST of each genome in the Datasets 1 and 2 and categorized them as “GC1” or “non-GC1” (see Materials and Methods: “MLST classification” section). Then we run the DBGWAS program using the Dataset 1 genome sequences as input; we obtained a total of 1,622,573 distinct unitigs that represented sequences of diverse length of the Dataset 1 taking into account the genomic variation (63). The length of the unitigs obtained was between 31 and 32,759 bp. We used the variant matrix built by DBGWAS to create the binary matrix used as input for the SVN algorithm. Unitigs that were found in less than 50 genomes from the Dataset 1 were discarded; we kept in the input binary matrix only the unitigs of size 31 to 385 bp.

Parameter tuning and model validations were performed using a 5-fold cross- validation and a grid search over a range of given values to determine the SVM kernel and hyperparameters that generated the best area under the curve (AUC). Once we obtained the SVM model that best fitted the Dataset 1 genomes, we extracted the first 100 unitigs that contributed most to *A. baumannii* GC1 strain prediction (Tables 1 and S3) according to the values of the features weight vector. By using blastn, we corroborated if the unitigs predicted by SVM were specific for *A. baumannii* GC1 detection. For this purpose, we searched for the unitig sequences in the genomes of the Dataset 2 to assess which were the unitigs with the greatest number of matches within each genome class (*A. baumannii* GC1 or non-GC1) (Tables 2 and S4). We also annotated the genome location and gene product related to the 100 unitigs (Tables 1, S3, 3 and S5).

**Table 1.**
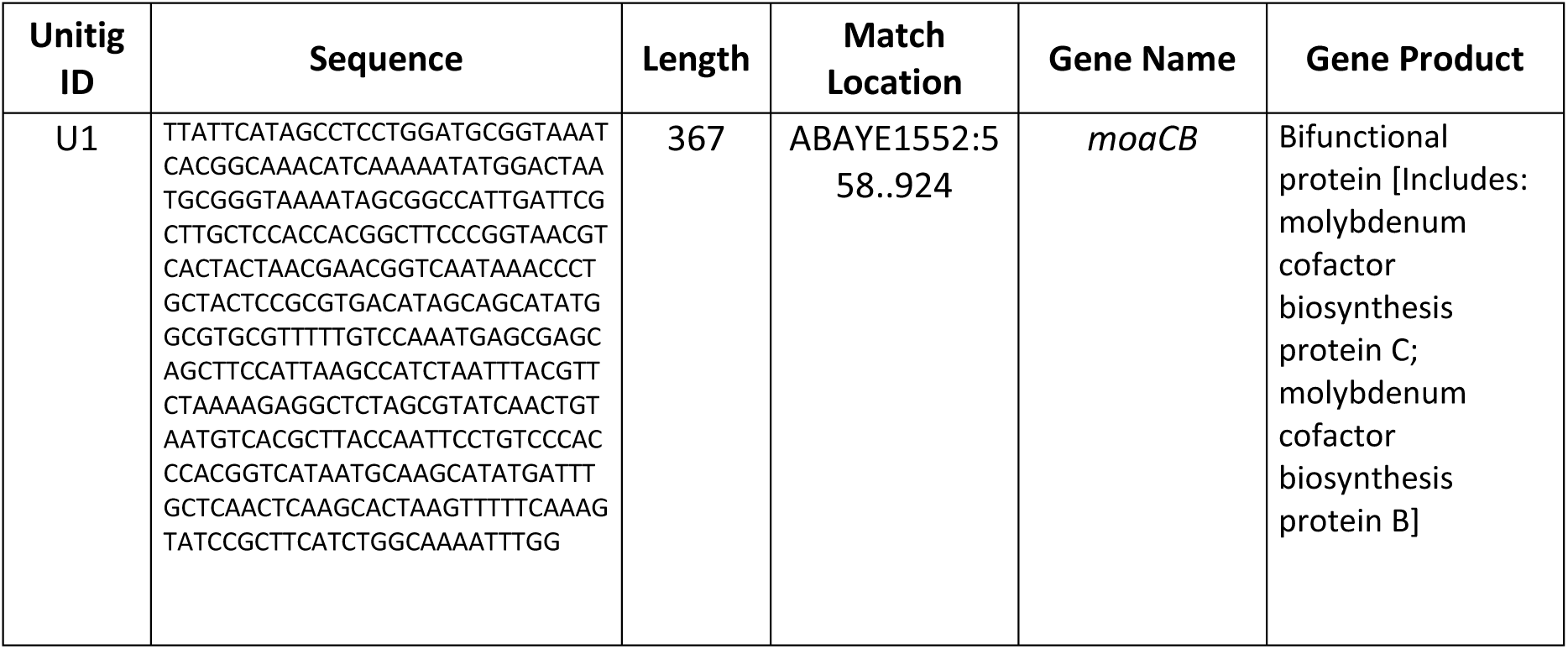
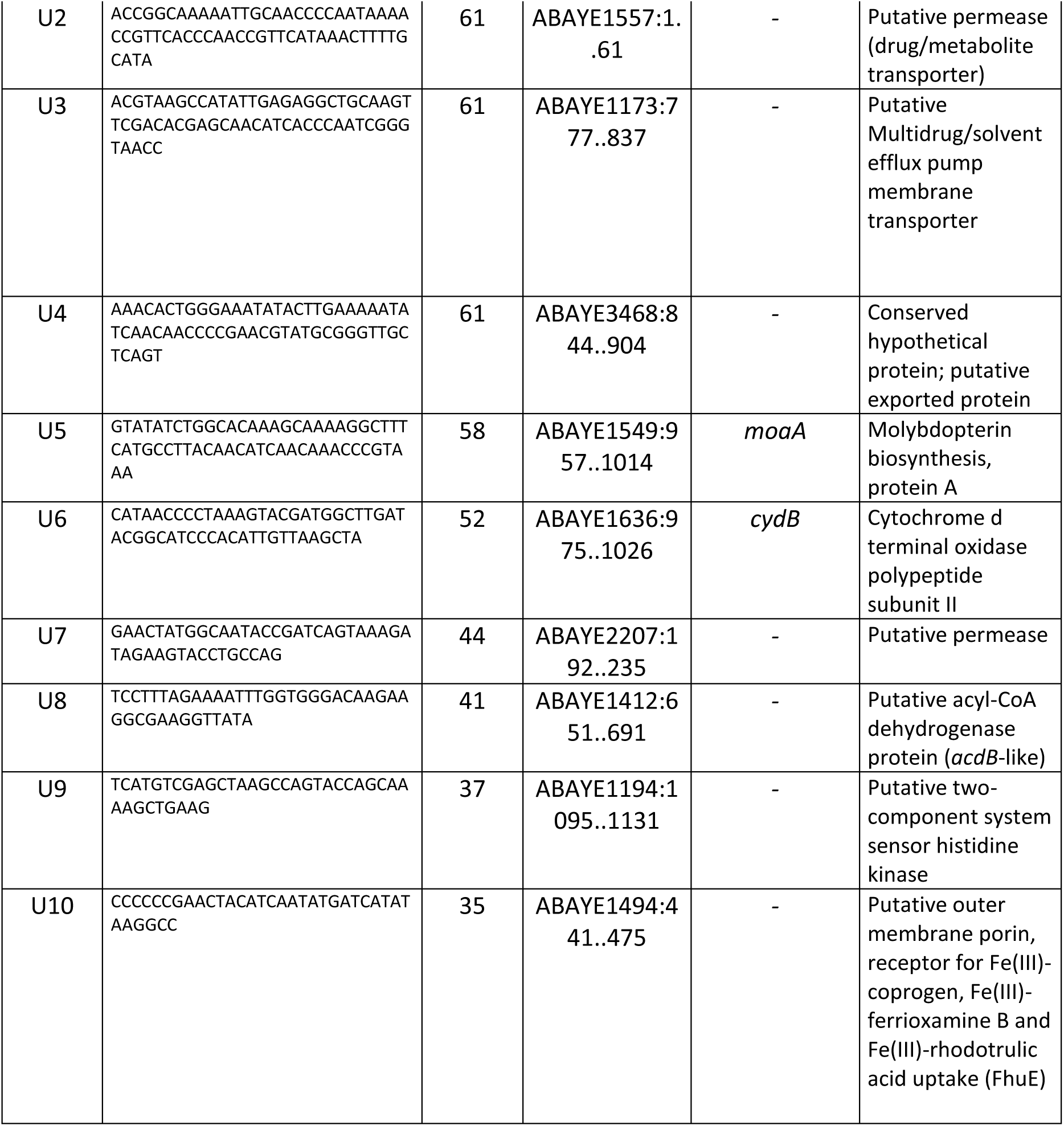
Putative *A. baumannii* GC1 biomarkers obtained by SVM. The table details the unitig ID, unitig sequence, unitig length, the location where the unitig sequence matched in *A. baumannii* AYE genome (AN:CU459141.1) or *A. baumannii* AB0057 genome (AN: CP001182.2), and the gene name corresponding to the genome region matched and the gene product of the first 10 unitigs that contributed most to *A. baumannii* GC1 genome prediction according to the values of the features weight vector. *A. baumannii* AYE strain genome was used as *A. baumannii* GC1 reference to locate the unitigs sequences. However, when the unitig was not found in *A. baumannii* AYE genome, *A. baumannii* AB0057 strain genome was used instead. The prefix “REGION” was used when the match occurred either in an intergenic region or in a combination of intergenic region and a gene. The complete list of the 100 unitigs predicted by SVM is detailed in Supplementary Table S3.

**Table 2.** Analysis of putative *A. baumannii* GC1 biomarkers obtained by SVM matches within the Dataset 1 and 2. . The table details the unitig ID of the first 10 unitigs that contributed most to *A. baumannii* GC1 genome prediction according to the values of the features weight vector, the number of *A. baumannii* GC1 and non-GC1 genomes typified by MLST that matched the unitig within the Dataset 1 and 2 using blastn and the p-values of Fisher’s exact test using a significance level of 0.05. Fisher’s exact test was calculated in R considering the nominal variables “Dataset Source” (Dataset 1 or Dataset 2) and “Matched” (Yes or No). The total number of *A. baumannii* GC1 and non-GC1 genomes that matched / not matched the unitigs in each dataset was used for calculation. Total data of the 100 unitigs predicted by SVM is detailed in Supplementary Table S4.

| Unitig ID | Dataset 1 |  | Dataset 2 |  | P-value (proportion of GC1 genome matches in the Dataset 1 not differs from proportion of GC1 matches in the Dataset 2) | P-value (proportion of non-GC1 genome matches in the Dataset 1 not differs from proportion of non-GC1 matches in the Dataset 2) |
| --- | --- | --- | --- | --- | --- | --- |
|  | Number of GC1 genomes matching the unitig | Number of non-GC1 genomes matching the unitig | Number of GC1 genomes matching the unitig | Number of non-GC1 genomes matching the unitig |  |  |
| U1 | 200/200 | 0/300 | 312/312 | 0/4487 | P = 1 | P = 1 |
| U2 | 200/200 | 0/300 | 311/312 | 0/4487 | P = 1 | P = 1 |
| U3 | 200/200 | 0/300 | 311/312 | 3/4487 | P = 1 | P = 1 |
| U4 | 200/200 | 0/300 | 310/312 | 43/4487 | P = 0.523 | P = 0.1106 |
| U5 | 200/200 | 0/300 | 311/312 | 0/4487 | P = 1 | P = 1 |
| U6 | 200/200 | 0/300 | 306/312 | 1/4487 | P = 0.0862 | P = 1 |
| U7 | 200/200 | 0/300 | 309/312 | 2/4487 | P = 0.2845 | P = 1 |
| U8 | 200/200 | 0/300 | 312/312 | 1/4487 | P = 1 | P = 1 |
| U9 | 200/200 | 0/300 | 309/312 | 5/4487 | P = 0.2845 | P = 1 |
| U10 | 200/200 | 0/300 | 311/312 | 0/4487 | P = 1 | P = 1 |

### Building SCM models for *A. baumannii* GC1 identification

In an additional approach, we applied the SCM algorithm implemented in Kover (30). We used as input the k-mers of the 500 *A. baumannii* genomes contained in the Dataset 1 considering the class label for each genome (*A. baumannii* GC1 or non-GC1). We ran Kover using k-mer sizes ranging from 31 to 127. Although output rules could be conjunctions (logical-AND) or disjunctions (logical-OR), we obtained 49 simple rules without conjunctions or disjunctions in the models of our study (Table 3 and S5). Rules depicted the presence (n=10) or the absence (n=39) of k-mers in the Dataset 1 genomes. Besides, we registered the number of matches of the k-mer sequences against *A. baumannii* GC1 and non-GC1 genomes contained in the Dataset 1 (Table 4 and S6); we also annotated the k-mer sequences (Tables 4 and S6).

**Table 3.**
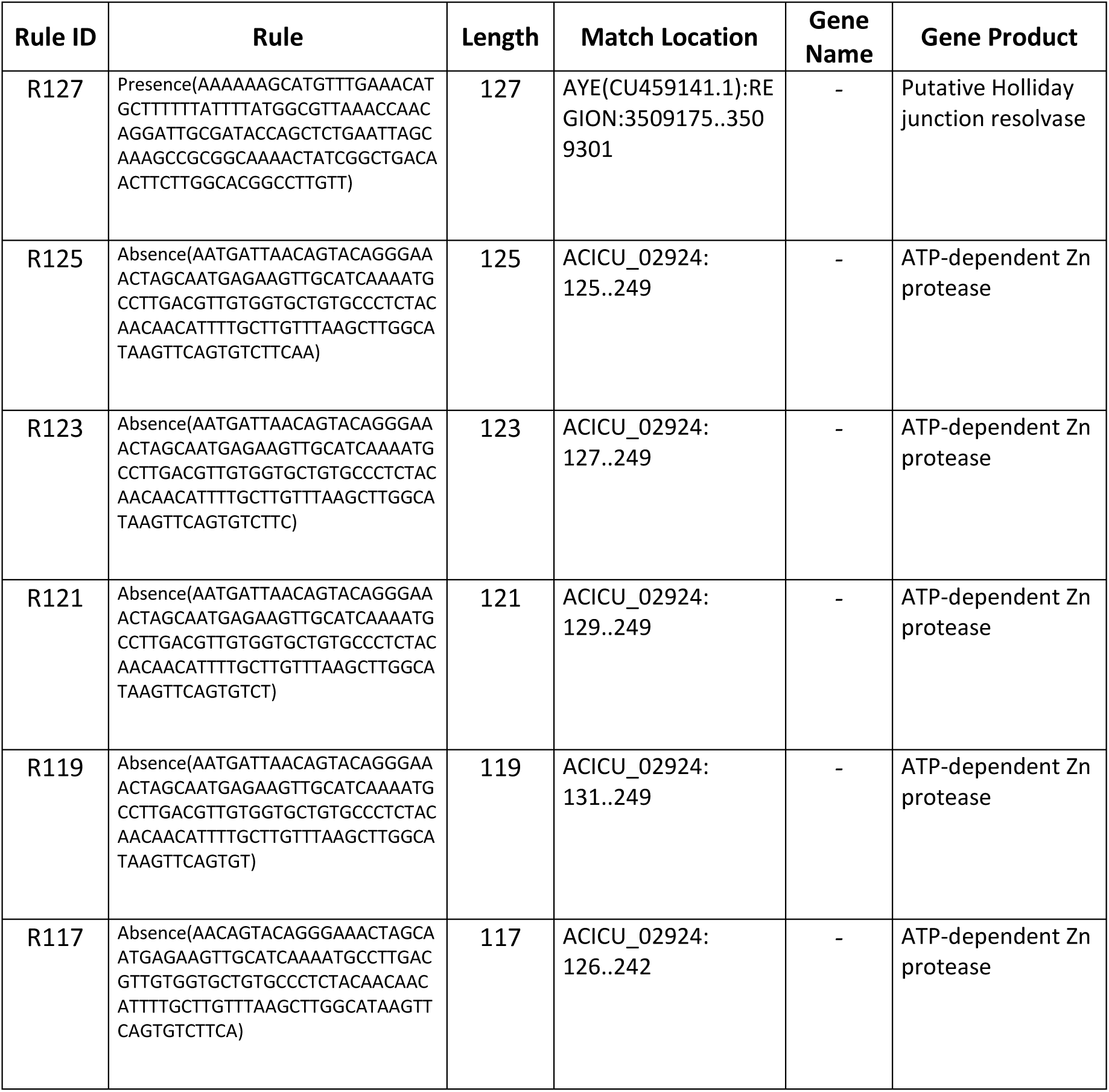
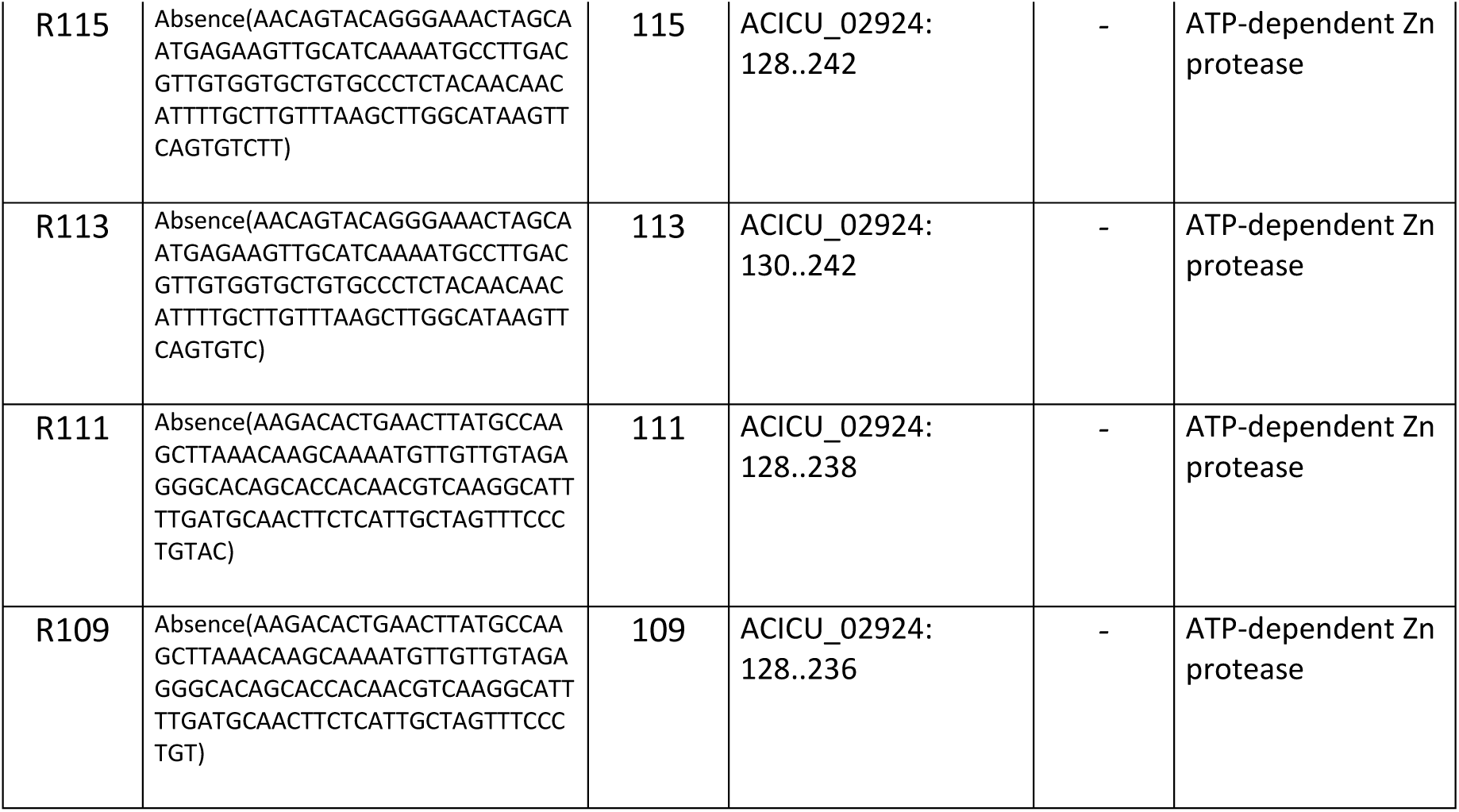
Rules obtained by the SCM models. The table details the rule ID, the rule output from the SCM model, k-mer length and the location where the rule sequence matched in *A. baumannii* AYE genome (AN:CU459141.1) or *A. baumannii* ACICU genome (AN: CP000863.1), and the gene name corresponding to the genome region matched and the gene product of the 10 larger k-mer sequences targeted by the SCM rules. *A. baumannii* AYE and ACICU genomes were used as *A. baumannii* GC1 and non-GC1 references, respectively, to locate the k-mer sequences. The prefix “REGION” was used when the match occurred either in an intergenic region or in a combination of intergenic region and a gene. The complete list of the 49 rules obtained by the SCM models is detailed in Supplementary Table S5.

**Table 4.** Analysis of the SCM rules matches within the Dataset 1 and 2. . The table details the rule ID of the 10 larger k-mer sequences targeted by the SCM rules, the number of *A. baumannii* GC1 and non-GC1 genomes typified by MLST that matched the rule within the Datasets 1 and 2 using blastn and the p-values of Fisher’s exact test using a significance level of 0.05. Fisher’s exact test was calculated in R considering the nominal variables “Dataset Source” (Dataset 1 or Dataset 2) and “Matched” (Yes or No). The total number of *A. baumannii* GC1 and non-GC1 genomes that matched / not matched the rules in each dataset was used for calculation. Total data of the 49 SCM rules is detailed in Supplementary Table S6.

| Rule ID | Dataset 1 |  | Dataset 2 |  | P-value (proportion of *A. baumannii* GC1 genome matches in the Dataset 1 not differs from proportion of GC1 matches in the Dataset 2) | P-value (proportion of *A. baumannii* non-GC1 genome matches in the Dataset 1 not differs from proportion of non-GC1 matches in the Dataset 2) |
| --- | --- | --- | --- | --- | --- | --- |
|  | Number of *A. baumannii* GC1 genomes matching the rule | Number of *A. baumannii* non-GC1 genomes matching the rule | Number of *A. baumannii* GC1 genomes matching the rule | Number of *A. baumannii* non-GC1 genomes matching the rule |  |  |
| R127 | 200/200 | 4/300 | 310/312 | 166/4487 | P = 0.523 | P = 0.03405 |
| R125 | 200/200 | 1/300 | 310/312 | 13/4487 | P = 0.523 | P = 0.5964 |
| R123 | 200/200 | 1/300 | 310/312 | 13/4487 | P = 0.523 | P = 0.5964 |
| R121 | 200/200 | 1/300 | 310/312 | 13/4487 | P = 0.523 | P = 0.5964 |
| R119 | 200/200 | 1/300 | 310/312 | 13/4487 | P = 0.523 | P = 0.5964 |
| R117 | 200/200 | 1/300 | 310/312 | 12/4487 | P = 0.523 | P = 0.5693 |
| R115 | 200/200 | 1/300 | 310/312 | 12/4487 | P = 0.523 | P = 0.5693 |
| R113 | 200/200 | 1/300 | 310/312 | 12/4487 | P = 0.523 | P = 0.5693 |
| R111 | 200/200 | 1/300 | 310/312 | 12/4487 | P = 0.523 | P = 0.5693 |
| R109 | 200/200 | 1/300 | 310/312 | 12/4487 | P = 0.523 | P = 0.5693 |

We observed that the rules obtained by the SCM models selected fragments of different length that matched the loci ACICU_02924 (n=23), ACICU_02095 (n=5), ABAYE3455 (n=5), ACICU_01506 (n=2) and ABAYE2468 (n=2) (Table 3 and S5). This fact could be caused by point mutations contained in the loci that the SCM models associated with *A. baumannii* GC1 prediction would have made as previously found (30).

### Selection of candidate biomarkers for rapid detection of *A. baumannii* GC1

We were interested in obtaining the longest DNA sequences shared by all *A. baumannii* GC1 genomes, and at the same time, without matches among *A. baumannii* non-GC1 genomes. For this purpose, unitigs and k-mer sequences obtained as candidate biomarkers for *A. baumannii* GC1 using the SVM and SCM algorithms were sorted in descending order according to the sequence length. Then, we considered the number of *A. baumannii* GC1 genomes that matched the unitig sequence or the SCM rule and sorted the candidate sequences in ascending order according to the number of *A. baumannii* non-GC1 genome matches.

First, we analyzed the SVM results. Data related to unitigs obtained are listed in Tables 1, 2, S3, S4 and, Figure 2. We named unitigs from U1 to U100. We observed that the unitigs named U1 to U12 were found in 100% of *A. baumannii* GC1 genomes while they were not found in *A. baumannii* non-GC1 genomes from the Dataset 1. However, U1 was the only unitig found in 100% of *A. baumannii* GC1 genomes within the Datasets 1 and 2, as well as absent in *A. baumannii* non-GC1 genomes of both datasets, being a specific biomarker for *A. baumannii* GC1 identification. Despite the U8 sequence was found in 100% of *A. baumannii* GC1 genomes within the Dataset 1 and 2, it was found in 1/4487 of *A. baumannii* non-GC1 genomes from the Dataset 2. The U8 sequence matched *A. baumannii* AYE genome in the locus tag ABAYE1412 between 651 and 691 coordinates (Table 1 and S3). The fragment is part of a gene that encodes a putative acyl-CoA dehydrogenase protein (*acdB*-like). As U8 sequence was not found exclusively in *A. baumannii* GC1 genomes, we discarded it as a possible biomarker of *A. baumannii* GC1. U4 was the sequence among these 12 unitigs that had the higher number of matches (43/4487) with *A. baumannii* non-GC1 genomes from the Dataset 2.

**Figure 2.**
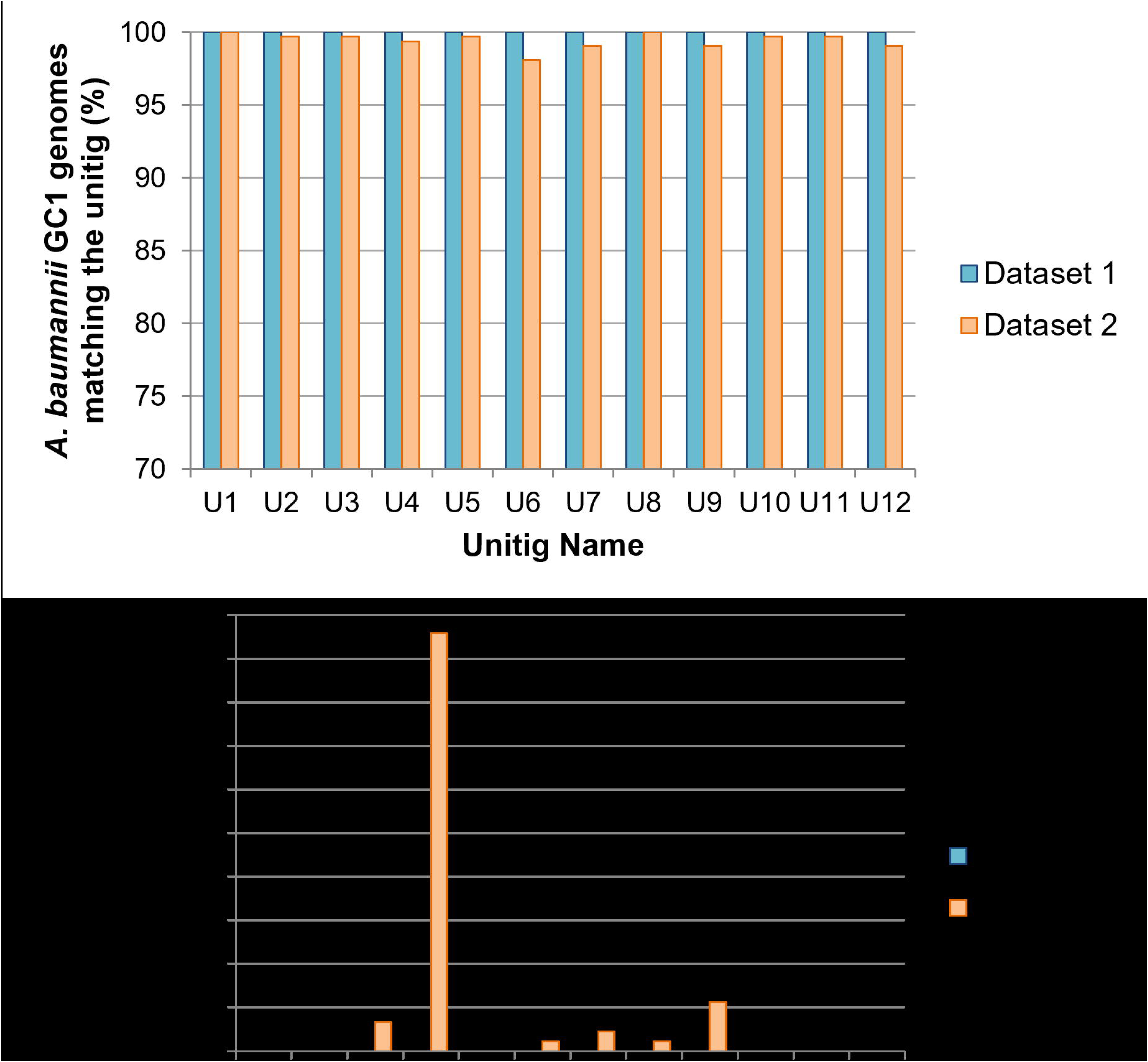
Percentage of genomes in the Dataset 1 and Dataset 2 matching unitig sequences U1 to U12. (A) *A. baumannii* GC1 genomes, (B) *A. baumannii* non-GC1 genomes.

The SCM results are listed in Tables 3, 4, S5, S6, and Figure 3. We observed that 43/49 rules targeted 100% of *A. baumannii* GC1 genomes from the Dataset 1 but also targeted between 1 and 4 of *A. baumannii* non-GC1 genomes from the Dataset 1. Within these 43 rules, two sets of rules targeted only 1 *A. baumannii* non-GC1 genome from the Dataset 1; as an example of this, 26/49 rules targeted genome with AN: GCF_000248195.1 (ST 69) and 1/49 rules (R49) targeted genome with AN: GCF_000453745.1 (ST 2) (Figure 3). Concerning the Dataset 2, 12/49 rules targeted 100% of *A. baumannii* GC1 genomes but also targeted several *A. baumannii* non-GC1 genomes from the Dataset 2. Since no rule obtained can uniquely identify *A. baumannii* GC1 genomes contained in our datasets, we were unable to obtain a putative biomarker from the results of the SCM models.

**Figure 3.**
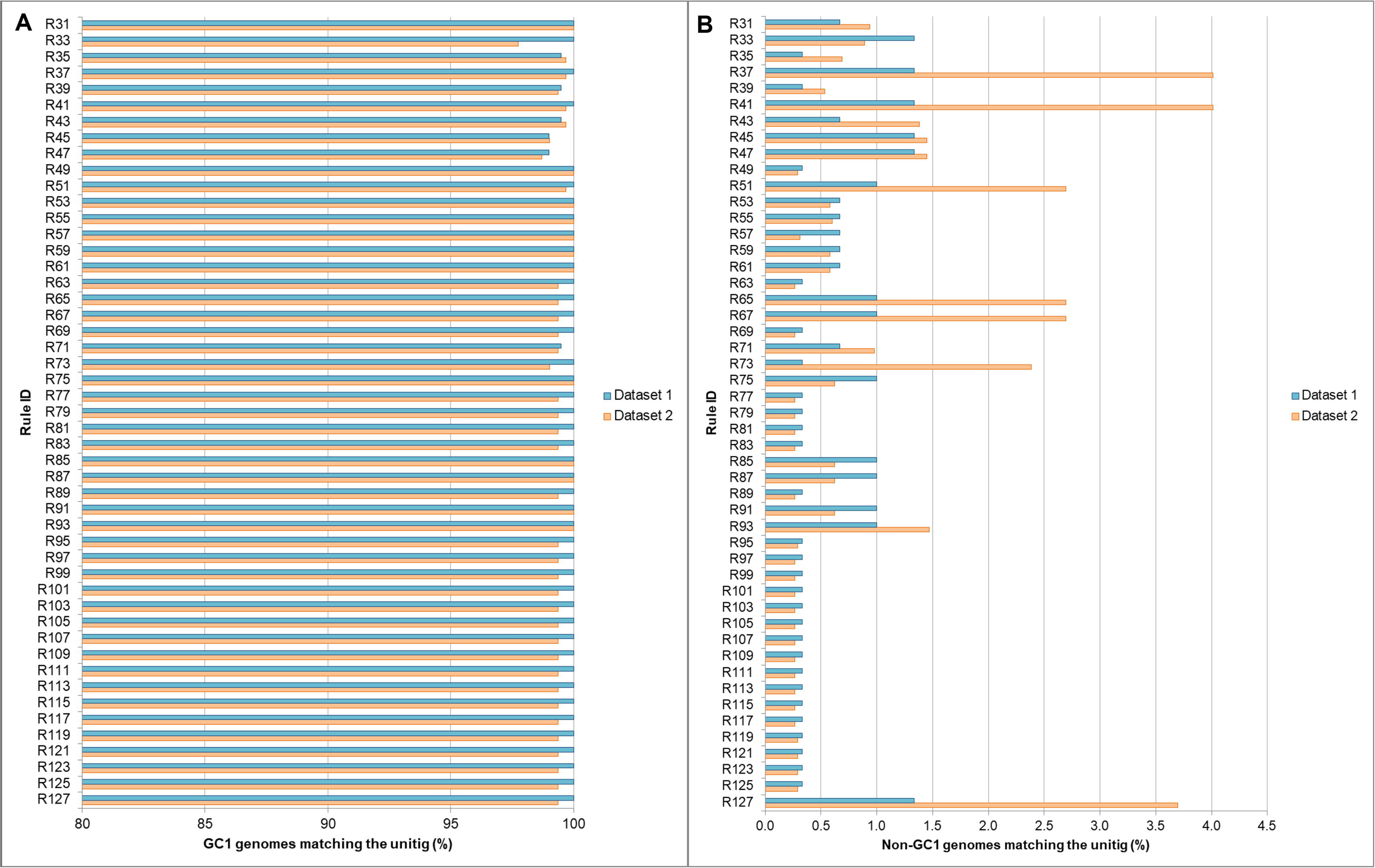
Percentage of genomes in the Dataset 1 and Dataset 2 matching the SCM rules. (A) *A. baumannii* GC1 genomes, **(B)** *A. baumannii* non-GC1 genomes. We observed that within the 43 rules which targeted 100% of *A. baumannii* GC1 genomes from the Dataset 1, two set of rules targeted only 1 non-GC1 from the same dataset. Rules R125, R123, R121, R119, R117, R115, R113, R111, R109, R107, R105, R103, R101, R99, R97, R95, R89, R83, R81, R79, R77, R73, R69, R63, R39 and, R35 targeted genome with AN: GCF_000248195.1 (ST 69) and R49 targeted genome with AN: GCF_000453745.1 (ST 2). Regarding the Dataset 2, rules R93, R91, R87, R85, R75, R61, 211 R59, R57, R55, R53, R49 and R31 targeted 100% of *A. baumannii* GC1 genomes but also targeted several *A. baumannii* non-GC1 genomes from the Dataset 2 (66/4487, 28/4487, 28/4487, 28/4487, 213 28/4487, 26/4487, 26/4487, 14/4487, 27/4487, 26/4487, 13/4487, 42/4487 genomes, respectively).

When using the Dataset 2 to validate the results obtained by the SVM and SCM models from the Dataset 1, we observed that in the case of the SVM classifier there was no statistically significant difference (P > 0.05) in the number of matches of the unitigs regardless of the genome dataset (Dataset 1 or 2) except for the unitig U69 (Tables 2 and S4). In the case of U69, we found a significant difference between the number of matches with *A. baumannii* non-GC1 genomes from the Dataset 1 and 2 (P = 0.01077). Concerning the SCM classifier, we observed that there was no statistically significant difference in the number of *A. baumannii* GC1 and non-GC1 genomes from the Dataset 1 and 2 that matched the rules (P > 0.05) (Tables 4 and S6). As we mentioned, the SVM model predicted the U1 sequence as a specific biomarker for *A. baumannii* GC1 genomes. The U1 sequence had 367 nucleotides and matched *A. baumannii* AYE genome in the locus tag ABAYE1552 between coordinates 558 and 924 (Table 1 and S3). This region corresponds to a fragment of the *moaCB* gene which encodes a bifunctional protein that includes the molybdenum cofactor biosynthesis protein C and protein B (64). In particular, the U1 sequence matched the region of the *moaCB* gene, which encodes for MoaB protein. The molybdenum cofactor (Moco) is an essential component of a large family of enzymes involved in carbon, nitrogen, and sulfur metabolism whose biosynthetic pathway is evolutionarily conserved. The MoaC protein, together with the MoaA protein, is involved in the first step of Moco biosynthesis (64). Interestingly, various studies have linked in-host survival of prevalent pathogenic bacteria such as *Mycobacterium tuberculosis*, *Escherichia coli*, and *Salmonella enterica* to the presence of functional molybdoenzymes (65–68). In our study, we found that the U1 sequence was 100% conserved in *A. baumannii* GC1 genomes (0 SNPs) while it was a variable region with 1 to 74 SNPs in the total of 4987 *A. baumannii* non-GC1 genomes from the Dataset 1 and 2 rendering 94 allelic variants in *A. baumannii* non-GC1 genomes. According to these results, the sequence of the *moaCB* gene comprised between 558 and 924 nucleotides allowed accurate discrimination for *A. baumannii* GC1 and non-GC1 genomes, becoming a biomarker that differentiates both groups. Moreover, we observed that the U1 sequence is conserved in all the *A. baumannii* GC1 genomes, but this is not the case for *A. baumannii* GC2. For this reason, U1 sequence could not be used as a biomarker for the early detection of *A. baumannii* GC2 strains.

Since 95 variants within 5299 *A. baumannii* strains were identified with multiple SNPs along the entire sequence of U1, we proposed that a strategy based on PCR amplification, would allow us to accurately differentiate *A. baumannii* GC1 from non-GC1 strains. For this reason, we designed the primeŕs pair BioM_GC1_ABA F 5’- TATTCATAGCCTCCTGGATGC-3’ and BioM_GC1_ABA R 5’-CCAGATGAAGCGGATACTTTG-3’, with coordinates 559 to 914 from ABAYE1552 locus tag representing 356 bp of U1 sequence (position 2 to 357). By a blastn search, we identified that this primer pair amplified only the U1 sequence recognized in *A. baumannii* GC1 strains amongst all the variants detected so far. The closest variants showed a mismatch of one nucleotide at the 3’ end of the reverse primer in *A. baumannii* ST163, ST411 and ST976 (AN GCF_015537765, GCF_000453725 and GCF_010500415, respectively).

Experimental analysis, with a total of 35 *A. baumannii* strains, which were 10 *A. baumannii* GC1 and 25 *A. baumannii* non-GC1 including 4 ST2 (GC2), 15 ST79, 5 ST119 and 1 ST404, was performed to test the primeŕs pair BioM_GC1_ABA F and BioM_GC1_ABA F (Figure S1). *A. baumannii* GC1 strains that were isolated more than 25 years apart were tested such as A144 (1997), A155 (1994) and HAX25Aba (2021) (69, 70). The pair of primers designed in this study only amplified by PCR the *A. baumannii* GC1 strains, and as expected, the subsequent DNA sequence analysis showed a 100% identity and query coverage with 356 bp of U1 as confirmed by the clustal alignment (Figure S2). Interestingly, *A. baumannii* Ab103 ST119 that has two mismatches in each primer, at positions 15 and 21 of the 21nts of BioM_GC1_ABA F, and at positions 9 and 21 of the 21nts of BioM_GC1_ABA R rendered also a negative result in this PCŔs conditions (see below). This result confirmed the specificity of this pair of primers.

The optimum PCR amplification was established in a final volume of 25 µl containing 0.625 U GoTaq® DNA Polymerase (cat. # M3005, Promega, USA), 5 µl 5 X Green GoTaq® buffer with MgCl_2_ for a final concentration of 1.5mM in the 1X reaction, 0.4 pM each dNTP, 1 µM each primer, and 5 µl of DNA from the boiling of 3-4 *A. baumannii* colonies in 100 µl of sterile H_2_O. DNA of *A. baumannii* GC1 strain A144 and water were used as positive and negative control, respectively. The PCR cycling conditions were an initial denaturation at 94 °C for five minutes, followed by 30 cycles of denaturation at 94 °C for 45 seconds, annealing at 58 °C for 45 seconds and extension at 72 °C for 30 seconds, with a final extension at 72 °C for five minutes. Alternatively, to this PCR to identify *A. baumannii* GC1, TaqMan assays could also provide a molecular typing method capable of being translated into clinical settings to differentiate *A. baumannii* GC1 and other relevant GC or ST.

### Evaluation of the performance of the SVM and SCM models

To avoid overfitting, a 5-fold cross-validation was performed on the training datasets used as input in both the SVM and SCM algorithms to select the best hyperparameter values with the highest AUC. Performance during testing was evaluated using the best hyperparameters obtained in terms of sensitivity, specificity, accuracy, precision, and F1 score (Tables 5, 6 and S7). The sensitivity of the SVM model was 1 ± 0.00, indicating that 100% of *A. baumannii* GC1 genomes were correctly identified within the testing dataset. Also, the SVM model achieved a high specificity (1 ± 0.00) when predicting *A. baumannii* non-GC1 genomes from the testing datasets. The precision value was 1 ± 0.00 when the model predicted that a genome was within *A. baumannii* GC1, being correct 100% of the time. Accuracy was 1 ± 0.00 indicating that 100% of *A. baumannii* GC1 and non-GC1 were correctly predicted. No false positives and no false negatives were predicted by the SVM model.

**Table 5.**
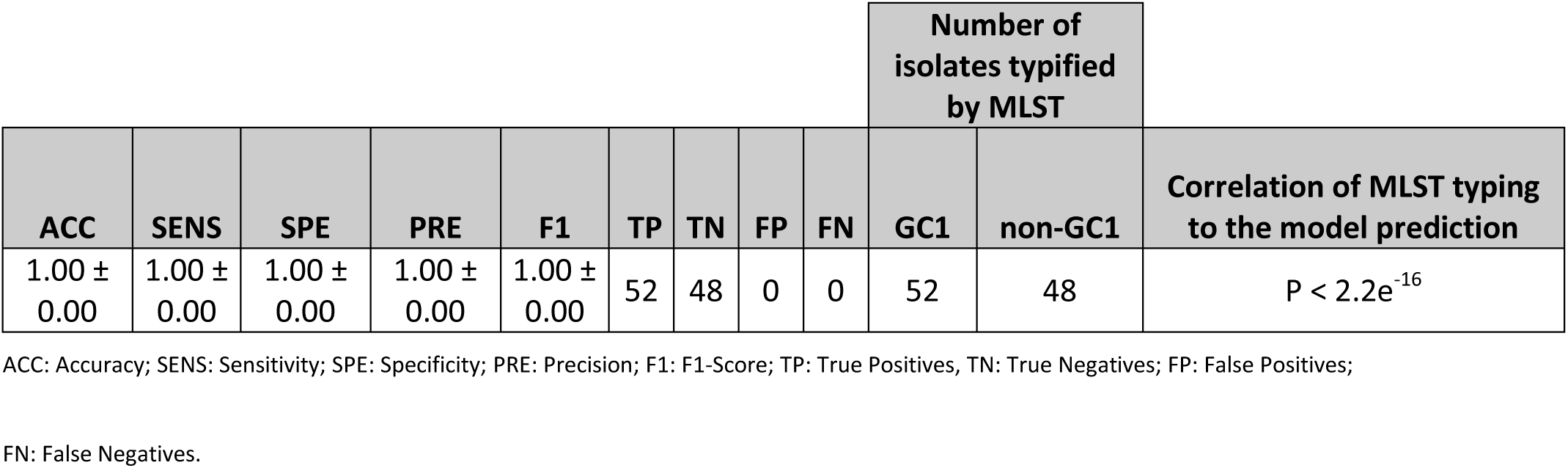
Prediction metrics on test dataset partitions from the Dataset 1 using the best performing SVM model. Correlation between MLST typing and model prediction was calculated using Fisher’s exact test in R level using a significance level of 0.05. The nominal variables “MLST Typing” and “Prediction” were considered during Fisher’s exact test calculation. The variable “MLST Typing” represented the genomes typed as *A. baumannii* GC1 (positive label) or non-GC1 (negative label) by MLST technique (true class). On the other hand, the variable “Prediction” represented the genomes predicted to be *A. baumannii* GC1 or non-GC1 by the SVM model (predicted class). We used the number of True Positives (TP), False Positives (FP), False Negatives (FN) and True Negatives (TN) in 2x2 contingency table. Null hypothesis used to evaluate the correlation between MLST typing and model prediction was: “True class (MLST Typing) and Predicted class are independent, knowing the value of one variable does not help to predict the value of the other variable”.

**Table 6.**
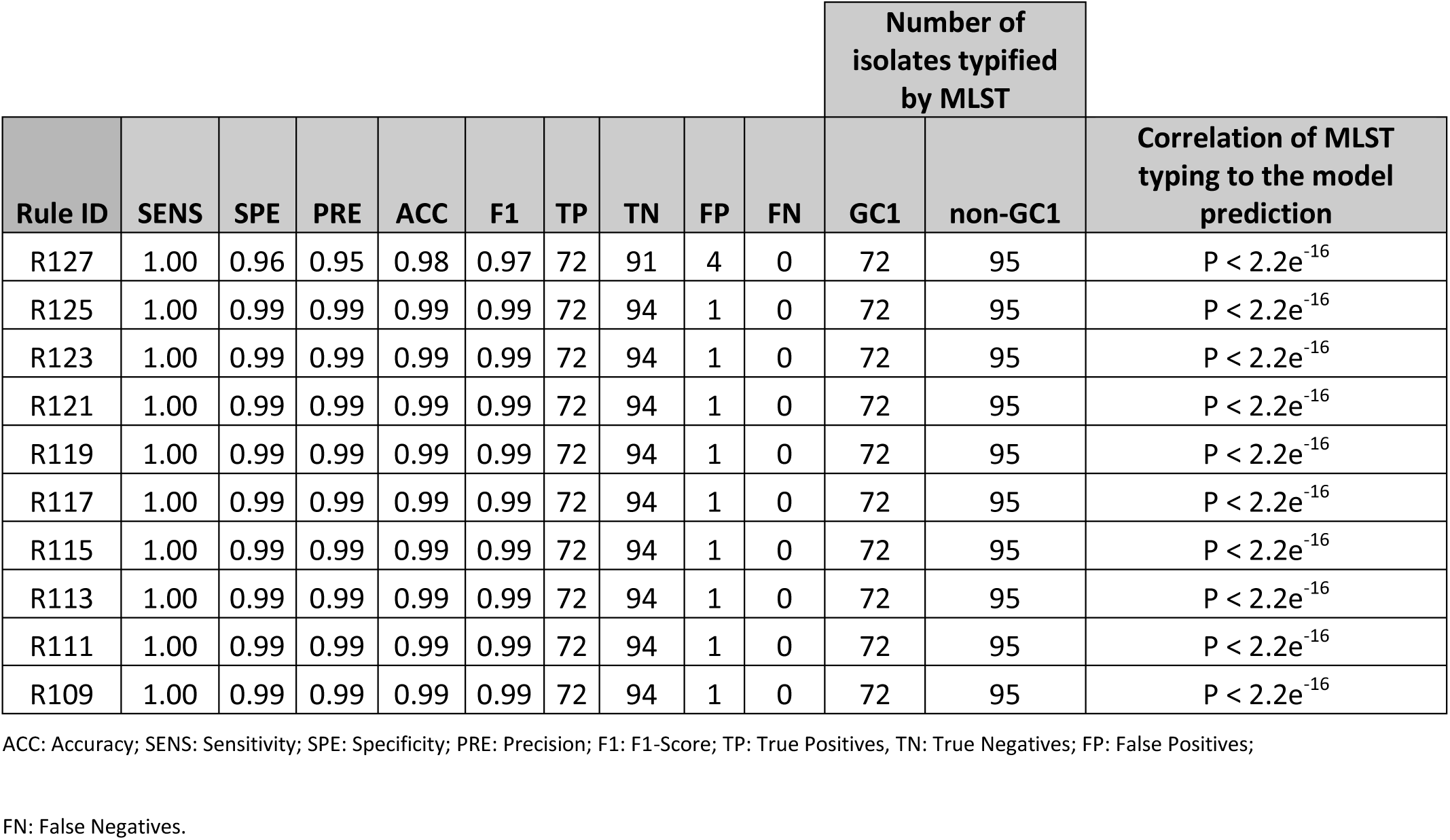
Prediction metrics on test dataset partitions from the Dataset 1 using the best performing SCM models. Correlation between MLST typing and model prediction was calculated using Fisher’s exact test in R using a significance level of 0.05. The nominal variables “MLST Typing” and “Prediction” were considered during Fisher’s exact test calculation. The variable “MLST Typing” represented the genomes typed as *A. baumannii* GC1 (positive label) or *A. baumannii* non-GC1 (negative label) by MLST technique (true class). On the other hand, the variable “Prediction” represented the genomes predicted to be *A. baumannii* GC1 or non-GC1 by the SCM model (predicted class). We used the number of True Positives (TP), False Positives (FP), False Negatives (FN) and True Negatives (TN) in 2x2 contingency table. Null hypothesis used to evaluate the correlation between MLST typing and model prediction was: “True class (MLST Typing) and Predicted class are independent, knowing the value of one variable does not help to predict the value of the other variable”. Total metrics of the 49 rules obtained by the SCM models is detailed in Supplementary Table S7.

|  |  |  |  |  |  |  |  |  |  | Number of isolates typified by MLST |  | Correlation of MLST typing to the model prediction |
| --- | --- | --- | --- | --- | --- | --- | --- | --- | --- | --- | --- | --- |
| Rule ID | SENS | SPE | PRE | ACC | F1 | TP | TN | FP | FN | GC1 | non-GC1 |  |
| R127 | 1.00 | 0.96 | 0.95 | 0.98 | 0.97 | 72 | 91 | 4 | 0 | 72 | 95 | $P < 2.2e^{-16}$ |
| R125 | 1.00 | 0.99 | 0.99 | 0.99 | 0.99 | 72 | 94 | 1 | 0 | 72 | 95 | $P < 2.2e^{-16}$ |
| R123 | 1.00 | 0.99 | 0.99 | 0.99 | 0.99 | 72 | 94 | 1 | 0 | 72 | 95 | $P < 2.2e^{-16}$ |
| R121 | 1.00 | 0.99 | 0.99 | 0.99 | 0.99 | 72 | 94 | 1 | 0 | 72 | 95 | $P < 2.2e^{-16}$ |
| R119 | 1.00 | 0.99 | 0.99 | 0.99 | 0.99 | 72 | 94 | 1 | 0 | 72 | 95 | $P < 2.2e^{-16}$ |
| R117 | 1.00 | 0.99 | 0.99 | 0.99 | 0.99 | 72 | 94 | 1 | 0 | 72 | 95 | $P < 2.2e^{-16}$ |
| R115 | 1.00 | 0.99 | 0.99 | 0.99 | 0.99 | 72 | 94 | 1 | 0 | 72 | 95 | $P < 2.2e^{-16}$ |
| R113 | 1.00 | 0.99 | 0.99 | 0.99 | 0.99 | 72 | 94 | 1 | 0 | 72 | 95 | $P < 2.2e^{-16}$ |
| R111 | 1.00 | 0.99 | 0.99 | 0.99 | 0.99 | 72 | 94 | 1 | 0 | 72 | 95 | $P < 2.2e^{-16}$ |
| R109 | 1.00 | 0.99 | 0.99 | 0.99 | 0.99 | 72 | 94 | 1 | 0 | 72 | 95 | $P < 2.2e^{-16}$ |

ACC: Accuracy; SENS: Sensitivity; SPE: Specificity; PRE: Precision; F1: F1-Score; TP: True Positives, TN: True Negatives; FP: False Positives;
FN: False Negatives.

Regarding the SCM models, the mean values of sensitivity, specificity, precision, and accuracy were 1 ± 0.01, 0.98 ± 0.01, 0.97 ± 0.01, and 0.99 ± 0.01, respectively. The mean rate of false positives was 1.11% while the mean rate of false negatives was 0.097%. All the rules obtained by the SCM models predicted false positives within the testing dataset (Table 6).

The training and the testing accuracy of the SVM and SCM models were above 0.99 (Figure 4), indicating that the SVM and SCM models, were not overfitted. The F1 score, which is the harmonic mean of precision and recall and is commonly used to compare different classification algorithms, was similarly high in the SVM and SCM models (1.00 and 0.99 ± 0.01, respectively). We used Fisher’s Exact Test to evaluate the performance of the SVM and SCM predictions comparing the actual genome classes (*A. baumannii* GC1 and non-GC1) typed by MLST and the predicted classes obtained by the models. In both cases, we obtained P < 2.2e-16 (Table 5, 6 and S7) indicating that the SVM and SCM models could significantly classify *A. baumannii* GC1 and non-GC1 strains. Despite these results, as the aim of this work was to find a biomarker that uniquely identifies *A. baumannii* GC1 genomes, the SVM model performed better than the SCM models since it did not predict false positives or false negatives.

**Figure 4.**
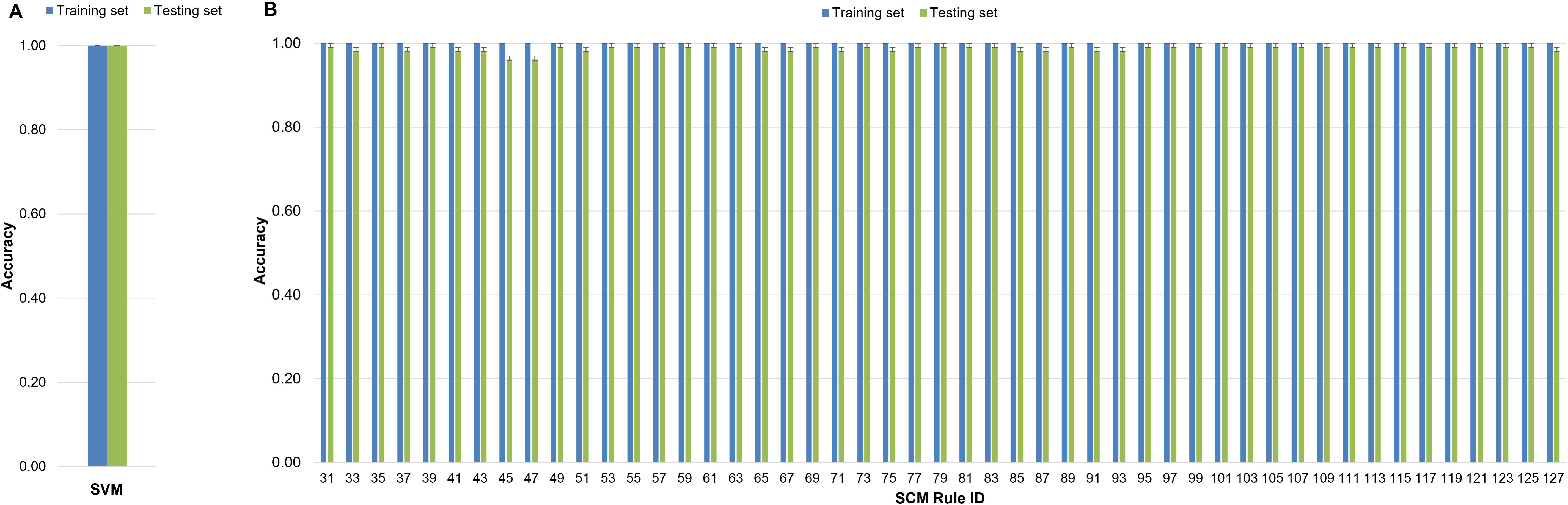
Mean Accuracy of the SVM and SCM models. Bars with blue color show the mean accuracy for the models on the training dataset. Bars with green color are the accuracy of the models on the test dataset. The models were run with the best hyperparameters selected from the 5-fold cross-validation. The error bars are standard deviations. **(A)** SVM model, **(B)** SCM models.

## DISCUSSION

High-risk clones or also named “superbugs” are dangerous clonal complexes with epidemic behavior equipped with exceptional resources both to infect the host and to evolve to extreme drug resistance phenotypes over time in the nosocomial niche (1, 71–77). The molecular understanding of the genetic and/or transcriptomic traits that lead to these capabilities remains still unknown (78, 79). Our study showed that ML applied to the study of high-risk clones can help not only in the identification of thoroughly accurate biomarkers but also contribute to disentangling molecular pathways that drive to epidemic lineages in the nosocomial niche which have not yet been completely deciphered. Accordingly, these findings could be used as therapeutic targets to reduce the dissemination of lineages with epidemic behavior. This is the case of the U1 sequence identified in the present study, which corresponded to 367 bp of the *moaCB* gene that encodes a bifunctional protein that includes the molybdenum cofactor biosynthesis protein C and protein B (64). Mononuclear molybdoenzymes (Mo enzymes) occur in organisms in all domains of life, where they mediate essential cellular functions such as energy generation and detoxification reactions (80). It has been shown that in bacterial pathogens, several processes such as the molybdate uptake, the cofactor biosynthesis, and the activities of Mo enzymes, affect fitness in the host as well as virulence (80). In addition to many studies on Mo enzymes that identified their crucial role in pathogenic species such as *E. coli*, *S. enterica*, *Campylobacter jejuni*, and *Mycobacterium tuberculosis*, some reports recently identified that these enzymes also contribute to the survival of ESKAPE pathogens (66, 81–85). Experimental studies must be undertaken to investigate the role of the *moaCB* gene in the virulence and fitness of *A. baumannii* which is also included in the ESKAPE group. Based on our results and previous experimental data in other pathogenic species (80), we can hypothesize that the U1 fragment of the *moaCB* gene in *A. baumannii* GC1 may be related to an essential metabolic pathway that play a vital role in the maintenance of epidemic clones in the hospital environment. Since it has been found that most of the Mo enzymes belong to groups that are unique to prokaryotes, these have been proposed as promising targets for the development of new antibiotic agents (80).

Given the variability observed in the biomarker U1 between *A. baumannii* GC1 and the 94 variants in *A. baumannii* non-GC1 strains, we propose a simple molecular biology strategy of one step of PCR amplification, to accurately differentiate *A. baumannii* GC1 from non-GC1 strains without performing MLST, or WGS and comparative genomics. The strategy proposed here, that was experimentally tested through this study, is accessible to a wide range of clinical and/or research laboratories. In addition, as the methodology consists of a single PCR, the detection of *A. baumannii* GC1 strains can be performed from the colony or directly from the clinical sample, giving the possibility of an early and simple diagnosis of this lineage.

Due to the increasing availability of bacterial WGS data, very active research has emerged on the use of this tool for genotype-phenotype prediction of antibiotic susceptibility (30, 33, 34, 44, 54, 86–88); however, there is no data available concerning the identification of high-risk clones based in WGS data excluding the MLST. Previous PCR-based studies used *bla*_OXA-51_ as one out of the three targeted genes to discriminate strains of *A. baumannii* GC1, GC2 and, GC3 (18, 19), and also the detection of deletion of 108 bp in the 5′-conserved segment (5′-CS) of the class 1 integron for identification of *A. baumannii* GC1 strains (19) have been proposed. Since *bla*_OXA-51_ has been found in other species (89–91) and the deletion of 108 bp as a biomarker differentiates partially two lineages within *A. baumannii* GC1 (7), and considering that both can be subjected to Lateral Genetic Transfer events, the analysis provided by ML in the present study supports a more accurate and solid implement to evaluate the presence of *A. baumannii* GC1 strains. Accordingly, U1 is part of an essential gene in *A. baumannii* GC1 genome (*moaCB*). This suggests that the optimization of metabolic genes from the core genome may be related to the exceptional abilities of high-risk clones. Interestingly, our data suggest that the accessory genome such as genomic islands or transposons involved in pathogenicity or antibiotic resistance, at least in *A. baumannii* GC1 strains, would not have played a causal role in the adaptation of this lineage to the hospital niche over time.

In our work, we applied two ML algorithms that differ substantially in methodology. The SVM algorithm used the number of occurring unitigs in *A. baumannii* GC1 and *A. baumannii* non-GC1 genomes to learn and predict clonal membership of the strains, while the SCM model used a greedy approach to construct conjunction or disjunction rules to find the most concise set of k-mers that allows for accurate *A. baumannii* GC1 or *A. baumannii* non-GC1 genome prediction. Previously, methods that combined the use of the SVM and SCM algorithms, and the representation of genomic data as k-mers were used to find genomic biomarkers to identify antibiotic resistance (30) or to predict antibiotic resistance from WGS data (54, 86). We ran the SCM algorithm through the Kover program with k-mer lengths between 31 and 127 nucleotides to be able to analyze all the possible rules obtained from these k-mers lengths. Also, we ran the SVM algorithm using unitigs that usually corresponded to a longer sequence than the individual equivalent k-mers. Unitigs are defined as the longest sequences that can be obtained when k-mers overlap by exactly k-1 nucleotides (63, 92, 93). In k-mer-based genome representations, the main downside is that the representation contains a lot of redundancy, since many k-mers are always present or absent simultaneously (e.g., gene deletion/ insertion). In this sense, it has been proposed to replace k-mers with unitigs (63, 92, 93).

We also proved by using statistical methods that the SVM and SCM models could significantly classify *A. baumannii* GC1 isolates (P < 0.05). While the SVM classifier predicted the U1 sequence as a specific biomarker for *A. baumannii* GC1 genomes, none of the rules obtained with the SCM models was able to uniquely identify *A. baumannii* GC1 genomes. All the rules obtained by the SCM models matched in *A. baumannii* GC1 and *A. baumannii* non-GC1 genomes from both the Dataset 1 and 2. Due to this result, it was not possible to obtain a sequence that could be used as a specific biomarker for *A. baumannii* GC1 strains from the rules obtained by the SCM models. A key step for the successful implementation of ML algorithms is the preparation of the input datasets (58, 94–96). In our study, we faced two issues related to the preparation of the Datasets 1 and 2. On one hand, a limitation in the program DBGWAS during the preparation of the matrix used in the SVM algorithm meant that only a total of 500 genomes could be included in the training dataset. Since our goal was to use the same training dataset for the SVM and SCM models, the 500 genomes were also used as input in the SCM algorithm. On the other hand, the scarcity of *A. baumannii* GC1 genomes available in GenBank compared to *A. baumannii* non-GC1 genomes caused the Dataset 2 to have visibly more genomes representing *A. baumannii* non-GC1 class. Despite this fact, the results obtained from the Dataset 1 and 2 remained consistent in 100% (P > 0.05) of the cases for the SVM model and more than 91.83% (P > 0.05) of the cases for the SCM models. Perhaps the limitation in the number of genomes in the training dataset is one of the reasons why the SCM models did not have enough samples to learn from *A. baumannii* GC1 genomes and therefore could not find a rule that uniquely identifies them. Conversely, the numbers of isolates predicted to be *A. baumannii* GC1 or non-GC1 by the SVM model using the unitig U1 were the same as the result obtained by the MLST typification technique. This fact indicated that the SVM model obtained was excellent in the classification of *A. baumannii* GC1 and non-GC1 genomes. It will be interesting to study in future works how the Dataset 1 splitting strategy (unitigs or k-mers), the number of genomes of each class (*A. baumannii* GC1 and non-GC1) in the Dataset 1, and the total number of *A. baumannii* genomes in the Dataset 1 impact on the SVM and SCM models predictions and performance. One possible approach could be using as input the binary matrix obtained from the unitigs representing the Dataset 1 genomes generated by the program DBGWAS in the program Kover and then to analyze the SCM model results. In the same way, we could obtain k-mer profiles (k-mer sizes ranging from 31 to 127) from the Dataset 1 genomes and the k-mer matrixes associated with each profile using the DSK k-mer counter (97). DSK is used by Kover to internally compute k-mer profiles from the input genomes (98). Then we could use the k-mer matrixes as input in the SVM algorithm. As result, we would obtain new putative sequence biomarkers from both approaches, and we could compare them with the ones obtained in the current work.

In conclusion, these results suggest that the application of ML to identify biomarkers for high-risk clones or superbugs can also be used at an exploratory level of great precision since can provide novel understandings about bacterial adaptation to the nosocomial niche. In turn, this data can contribute to the experimental work with the possibility to be further translated to the clinical settings. The SVM algorithm made genetic predictions based on the presence or absence of short genomic sequences present in *A. baumannii* GC1 and non-GC1 genomes, that detected a biomarker, U1, not related to Lateral Genetic Transfer neither to accessory genome nor to antibiotic pressures that uniquely identifies the CG1 strains, which in agreement with previous experimental works on the group of Mo enzymes, showed that the application of ML could be a powerful tool to discover new therapeutic targets for the development of new antibiotic agents.

## MATERIALS AND METHODS

### MLST classification

Multilocus sequence typing (MLST) of all genomes in the data collection was performed *in silico* using the mlst software developed by T. Seemann (https://github.com/tseemann/mlst) and the Pasteur’s MLST database and schema for *A. baumannii* (https://pubmlst.org/organisms/acinetobacter-baumannii). *A. baumannii* sequence type (ST) numbers ST 1, 19, 20, 81, 94, 328, 460, 623, 315, 717 and 1106 were classified into *A. baumannii* GC1 and others ST into *A. baumannii* non-GC1 genomes (Tables S1 and S2).

### Data collection

To perform an accurate ML analysis to identify an *A. baumannii* GC1 biomarker, we defined two datasets to do our studies.

The Dataset 1 was composed of 200 *A. baumannii* GC1 and 300 *A. baumannii* non- GC1 genomes obtained from Genbank and typified by MLST as previously described. These genomes were retrieved from the GenBank assembly database filtering by *Acinetobacter baumannii* in the search by organism option (https://www.ncbi.nlm.nih.gov/assembly/organism/, last accessed in July 2021). *A. baumannii* GC1 genomes included 18 genomes used in a previous study (Álvarez et al., 2020) and 182 *A. baumannii* genomes as scaffolds and contigs (Table S1). *A. baumannii* non-GC1 genomes included five genomes belonging to other high-risk epidemic clones such as ACICU (AN: CP031380.1) as representative of GC2, Naval-13 (AN: AMDR01000001.1) as representative of GC3, AB33405 (NZ_JPXZ00000000.1) as representative of local epidemic clone CC113 and both ATCC 17978 (AN: CP018664.1), and A118 (AN: AEOW01000000) as sporadic clones. We also included 205 *A. baumannii* non- GC1 genomes as scaffolds and contigs (Table S1). The Dataset 2 was composed of 312 *A. baumannii* GC1 and 4487 *A. baumannii* non-GC1 genomes (Table S2) retrieved from the GenBank assembly database (last accessed in July 2021). The number of genomes in this dataset was limited by *A. baumannii* GC1 and non-GC1 genomes available in the GenBank database at the time of the query. We used the Dataset 2 to validate using blastn searches the results obtained by the SVM model. (Table S2). STs were numbered according to the Pasteur scheme for MLST (Table S1 and S2).

### Machine learning analyses

The SVM classifier is based on the maximization of the margin around the hyperplane (*w^T^x + b*) separating samples or instances of the different classes (57, 99). Each instance *i* = 1,…,*m* consists of an N-dimensional feature vector *x_i_* and a class label *y_i_* ∈ {+1, −1}. The maximization of the margin corresponds to the following minimization:

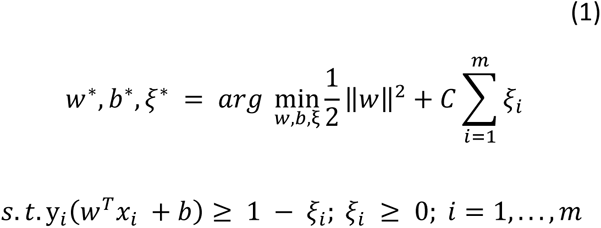

In this soft-margin SVM equation, *ξ_i_* is a penalty for misclassification or classification within the margin. Parameter *C* sets the weight of this penalty. The resulting weight vector *w* encodes the contributions of all features to the classifier (57).

We created a Python script using scikit-learn (100) to run the SVM algorithm. The script evaluated the classifier through a 5-fold cross-validation. In detail, the data was split into five consecutive folds (without shuffling) using the Python scikit-learn (sklearn) KFold function (https://scikit-learn.org/stable/modules/cross_validation.html#k-fold), and five models were built. Each fold was used once as a test set while the four remaining folds formed the training set. During each of the five iterations, the hyperparameter tuning was done using a 5-fold cross-validated grid search using the GridSearchCV function implemented in the sklearn.model_selection package (https://scikit-learn.org/stable/modules/generated/sklearn.model_selection.GridSearchCV.html#sklearn.model_selection.GridSearchCV) to find the best hyperparameters. We evaluated linear, polynomial (with a default degree of 3), radial basis function (RBF), and sigmoid kernels. We considered values between 0.01 and 100 for the penalty parameter of the error term (C) and values between 0.000001 and 10 for the gamma parameter. The predictions of all five iterations were compared using the AUC score (https://scikit-learn.org/stable/modules/model_evaluation.html#roc-metrics). Finally, we built up the classifier from the entire training set using the best hyperparameters (with the highest AUC) identified through cross-validation and applied the best model to the test set. The highest AUC were obtained using kernel = linear, gamma = 0.000001 and C = 0.01 (code available on GitHub at https://github.com/vealvarez/SVM_GC1).

The SCM (42), is a learning algorithm that produces models that are conjunctions (logical-AND) or disjunctions (logical-OR) of boolean-valued rules *r* : ℝ^*d*^ → {0,1}. Let us use *h(x)* to denote the output of model *h* on genome *x*. When *h* consists of a conjunction (i.e., a logical-AND) of a set of rules we have

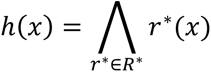

Whereas, for a disjunction (i.e., a logical-OR) of rules, we have

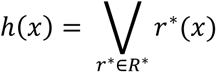

Given a set *R* of candidate rules, the SCM algorithm attempts to find a model that minimizes the empirical error rate 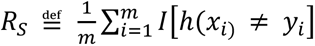, where *I*[True] = 1 and 0 otherwise while using the smallest number of rules in *R* (101).

We used the program Kover which implements the SCM algorithm (30, 98). Kover combines the SCM algorithm with the k-mer representation of genomes, which reveals uncharacteristically sparse models that explicitly highlight the relationship between genomic variations and the phenotype of interest (30). We ran the program using the Dataset 1 (see Input dataset preparation below) for k-mer sizes from 31 to 127 (taking only the odd numbers between them). The smallest value of k was set to 31 since extensive testing has shown that this size is optimal for bacterial genome assembly and has been employed for studies based on reference-free bacterial genome comparisons (92, 102). The greatest k-mer size was set to 127 since it is the maximum value accepted as a parameter in Kover. We chose only odd values of k to avoid the formation of palindromes (103). For each k-mer size, we split the Dataset 1 into a training dataset (2/3 of the genomes) and a testing dataset (1/3 of the genomes). Then, we trained the 49 models corresponding to each k-mer using a conjunction/disjunction model type. The best conjunctive and/or disjunctive model for each k-mer was selected using 5-fold cross-validation to determine the optimal rule scoring function with default parameters.

### Input dataset preparation for ML models

We split the 500 genomes included in the Dataset 1 into unitigs by using the program DBGWAS (63). Unitigs are stretches of DNA shared by the strains in a dataset. The DBGWAS method proposes connecting the overlaps of k-mers in a compressed de Bruijn graph (DBG) so that k-mers are extended using the adjacent sequence information in the population, forming unitigs present in the same set of samples as their constituent k-mers. During the first step of the DBGWAS process, the program built a variant matrix, where each variant is a pattern of the presence/absence of unitigs in each genome present in the Dataset 1 (63). We wrote a Python script (code available on GitHub at https://github.com/vealvarez/SVM_GC1) to format the variant matrix and create a presence/absence (coded with the values 1/0 respectively) binary matrix with unitigs as columns (features) and the accession numbers of the genomes as rows (instances). We used the binary matrix as input for the SVM algorithm. In this matrix, we discarded the data about low frequency unitigs (unitigs found in less than 50 *A. baumannii* GC1 and non-GC1 genomes). We also integrated the MLST data corresponding to the Dataset 1 genomes and created a two-column matrix used as input for the SVM algorithm. The first column of the matrix contained the accession number of the genomes, and the second column contained a binary variable that indicated whether each genome was typed or not as *A. baumannii* GC1 (-1 = *A. baumannii* non-GC1 genome / 1 = *A. baumannii* GC1 genome) according to MLST typing.

For the SCM approach, we first packaged the Dataset 1 sequences stored in FASTA files into a Kover dataset using the create from-contigs command. This command also received a tab-separated value (TSV) with the Dataset 1 genome classification according to MLST typing (*A. baumannii* GC1 or non-GC1). The first column of the TSV file described the genome accession number and the second column had the value 1 whether the genome belonged to *A. baumannii* GC1 or the value 0 otherwise. We ran Kover for k-mer sizes from 31 to 127. For each k-mer size, Kover constructed a reference-free input matrix based on k-mer profiles generated with the DSK k-mer counting software. A k-mer presence/absence binary matrix based on the Dataset 1 genomes was then created and used as input for the SCM models of each k-mer size.

### Unitig selection using SVM for putative biomarkers analysis

After obtaining the SVM model with the best hyperparameters, the values of the features weight vector (referred to as the hyperplane normal vector *w* in the formula (1)) were accessed through the attribute sklearn.model_selection.GridSearchCV.best_estimator.coef_. The values were sorted from highest to lowest and used to decide the relevance of each unitig sequence (associated with each weight value) during the model prediction (104, 105). It is worth mentioning that in our study, sequence unitigs were used as features in the models. The positive sign of a feature weight value indicates that the feature contributes to *A. baumannii* GC1 class prediction (represented by the value 1) and the negative sign indicates that the feature contributes to *A. baumannii* non-GC1 class (represented by the value -1) prediction (106). Considering the above mentioned, 100 unitig sequences with the highest weight values were selected to be analyzed as putative biomarkers of *A. baumannii* GC1 genomes.

### Machine learning performance metrics

The performance of the SVM and SCM models was evaluated in terms of sensitivity, specificity, accuracy, precision, and F1-score. They were defined as: sensitivity = TP/(TP + FN), specificity = TN/(TN + FP), accuracy = (TP + TN)/(TP + FP + TN + FN), precision = TP/ (TP + FP) and F1 score = 2 * (precision * sensitivity)/(precision + sensitivity). Where TP (true positives) was the number of *A. baumannii* GC1 strains predicted to be *A. baumannii* GC1, TN (true negatives) was the number of *A. baumannii* non-GC1 strains predicted to be *A. baumannii* non-GC1, FP (false positives) was the number of *A. baumannii* non-GC1 strains predicted to be *A. baumannii* GC1, and FN (false negatives) was the number of *A. baumannii* GC1 strains predicted to be *A. baumannii* non-GC1.

### Blastn searches

Blastn searches (107) were done using the Dataset 1 and Dataset 2 as subjects, and unitigs/k-mers that contributed most to *A. baumannii* GC1 genome prediction according to the SVM/SCM models were used as queries. We identified whether unitigs/k-mers matched known genes or intergenic regions and provide their putative function when possible. AYE (AN: CU459141.1) and ACICU (CP000863.1) genomes were used as a reference to annotate the genome location and gene product related to *A. baumannii* GC1 and non-GC1 genomes, respectively. In case the unitig was not found in the AYE genome, AB0057 (CP001182.2) genome was used instead. Also, we counted the number of *A. baumannii* GC1 and non-GC1 genomes matched by each unitig. We considered a cut-off E-value of E^−10^, 100% of identity, and 100% of query cover. To analyze the target of the primers designed below, Blastn searches (107) were done by using primer’s sequence as query and the Nucleotide collection (nr/nt) database of GenBank, or the Dataset 1 and Dataset 2 as the subject. Finally, Blastn searches (107) were also done by using the fragment of the U1 genomic biomarker of *A. baumannii* GC1 amplified with those primers (excluding the sequence of the primers) as query, and the Dataset 1 and Dataset 2 as the subject.

### Primer design

The primers to amplify the U1 genomic biomarker of *A. baumannii* GC1 were designed by using the Oligo primer analysis software version 6.22 (108, 109).

## SUPPLEMENTARY MATERIAL

Supplemental material is available online only.

## ACKNOWLEDGMENTS

The authors’ work was supported by the grant PUE 2522 from CONICET given to IMPaM.

We are grateful to the technical assistance of Dr. Gabriela Camicia and Nicolás Donis from CONICET.

DC, VA, and PQ contributed to the conception and design of the study. VA performed the script development. VA and PQ analyzed the results. DC and VA structured the work, wrote, and coordinated the drafts of the manuscript and did the final edition. VA wrote the results section. All authors contributed to the manuscript revision, read, and approved the submitted version.

## SUPPLEMENTAL FIGURES LEGENDS

**Figure S1.**
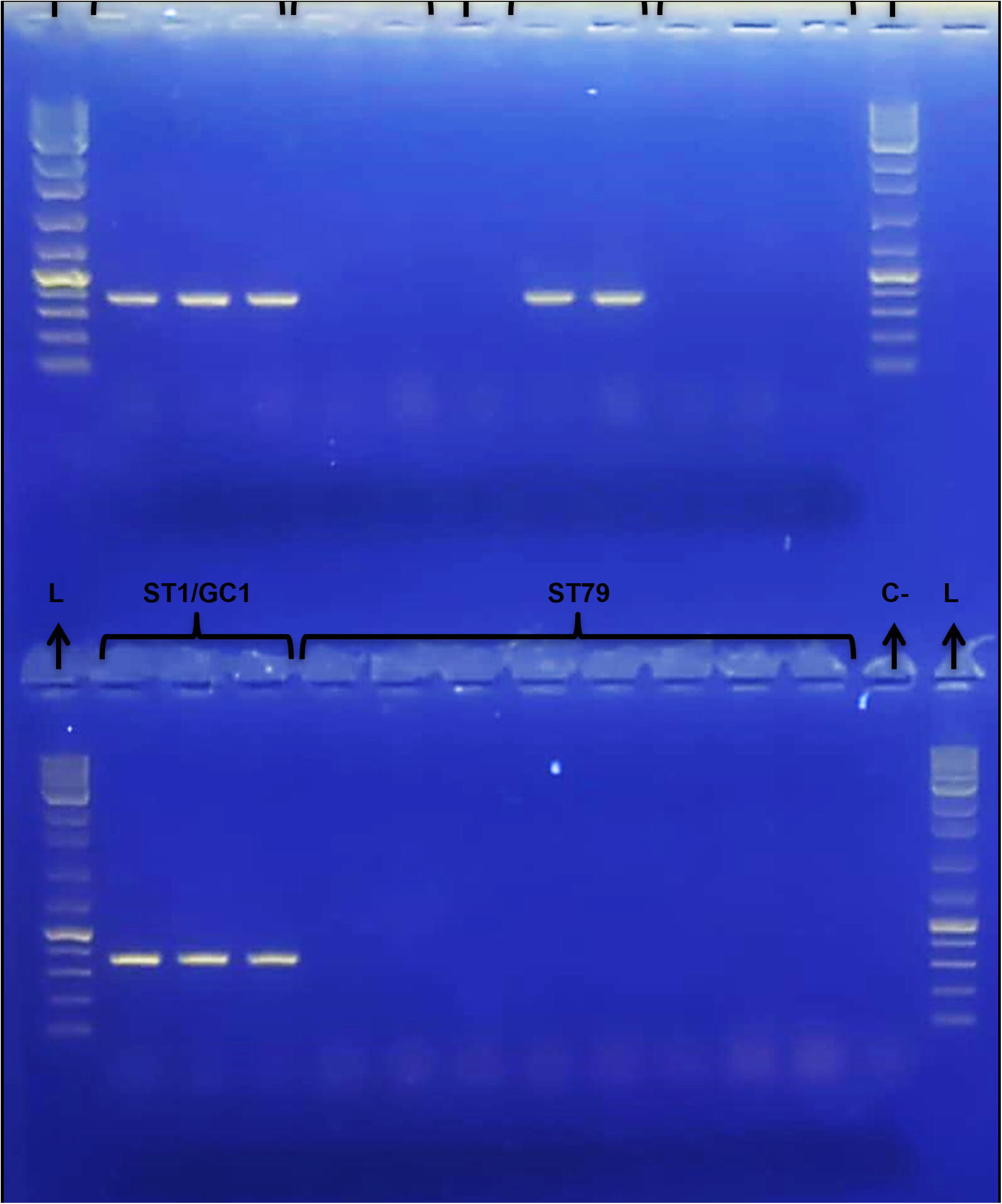
*A. baumannii* GC1 biomarker identification by PCR. PCR amplification of several *A. baumannii* strains is shown. Samples order at the upper wells correspond to Ladder, A144 ST1, A155 ST1, HAX19AbA ST1, HAX26Aba ST2, HAX28Aba ST2, A118 ST404, A110 ST1, A185 ST1, A325 ST119, A376 ST119, A103 ST119, ladder. Samples order at the lower wells correspond to Ladder, HAX22Aba ST1, HAX25Aba ST1, HAX27Aba ST1, Ab33405 ST79, Ab66 ST79, 186 ST79, F33943 ST79, A171 ST79, A177 ST79, A182 ST79, A384 ST79, negative control, ladder. Agarose gel 1.5%, electrophoresis in TAE buffer at 80V during 45 min. 1 Kbp DNA ladder (#K0178, Inbio Highway, Argentina).

**Figure S2.**
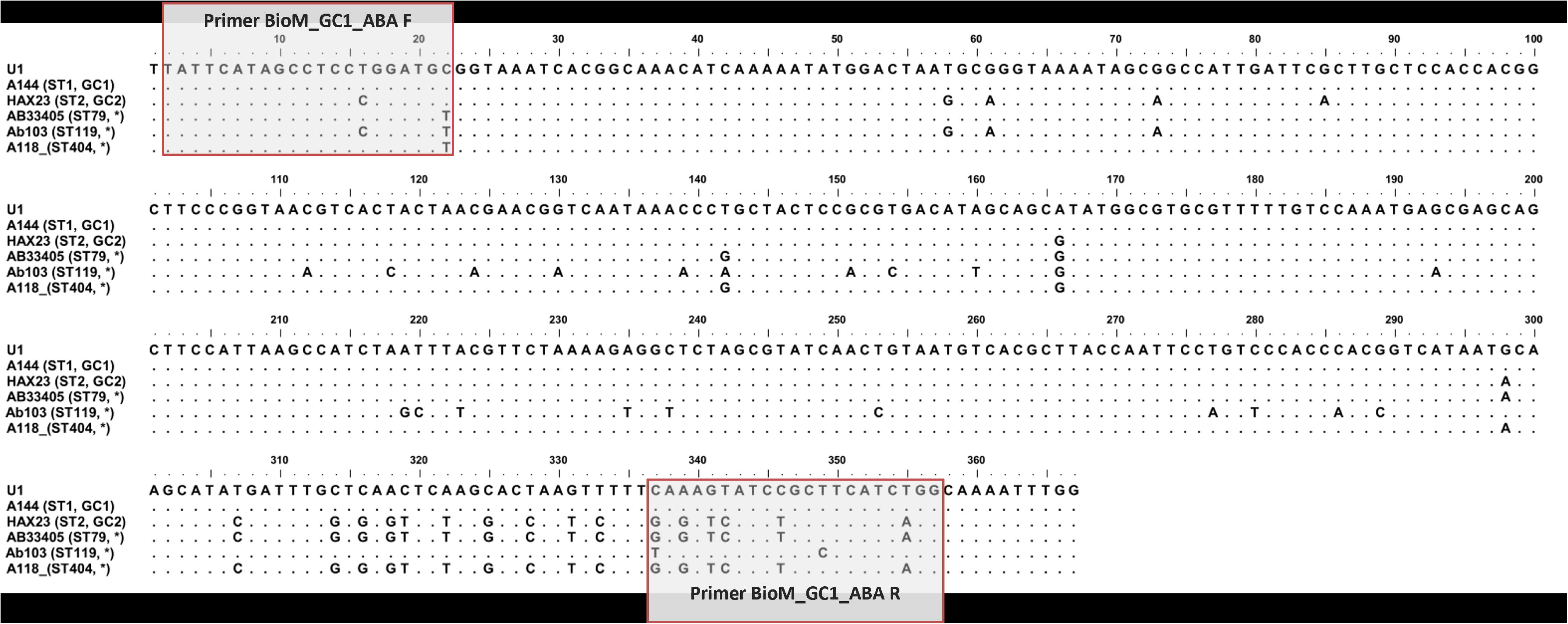
Alignment of the DNA sequence of U1 identified in *A. baumannii* strains tested by PCR. The primers pair used in the PCR are shown, which targets coordinates 559 to 914 from ABAYE1552 locus tag representing positions 2 to 357 of the 367 bp length U1 sequence identified in *A. baumannii* GC1 amplified by PCR. Sequence type (ST) according to Pasteur’s MLST schemes and Global Clones (GCs) are indicated in brackets. *The ST does not belong to a GC.

## SUPPLEMENTAL TABLES LEGENDS

**Table S1. *A. baumannii* genomes used as input dataset for SVM algorithm within the Dataset 1 genome collection.** GenBank accession number and classification of the 500 genomes in ML analysis. Genomes were classified as *A. baumannii* GC1 and non-GC1 according MLST Pasteur scheme.

**Table S2. *A. baumannii* genomes used as input dataset for SVM algorithm within the Dataset 2 genome collection**. Genomes were classified as *A. baumannii* GC1 and non-GC1 according MLST Pasteur scheme. ST Pasteur column with “-” value indicates that the genome does not yet have a ST assigned in the PubMLST database.

**Table S3. Putative GC1 biomarkers obtained by SVM.** The table details the unitig ID, unitig sequence, unitig length, the location where the unitig sequence matched in *A. baumannii* AYE genome (AN:CU459141.1) or *A. baumannii* AB0057 genome (AN: CP001182.2) and the gene name corresponding to the genome region matched and the gene product. *A. baumannii* AYE strain genome was used as *A. baumannii* GC1 reference to locate the unitigs sequences. However, when the unitig was not found in the AYE genome, *A. baumannii* AB0057 strain genome was used instead. The prefix “REGION” was used when the match occurred either in an intergenic region or in a combination of intergenic region and a gene.

**Table S4. Analysis of putative GC1 biomarkers obtained by SVM matches within the Dataset 1 and 2.** The table details the unitig ID of the first 100 unitigs that contributed most to *A. baumannii* GC1 genome prediction according to the values of the features weight vector, the number of *A. baumannii* GC1 and non-GC1 genomes typified by MLST that matched the unitig within the Dataset 1 and 2 using blastn and the p-values of Fisher’s exact test using a significance level of 0.05. Fisher’s exact test was calculated in R considering the nominal variables “Dataset Source” (Dataset 1 or Dataset 2) and “Matched” (Yes or No). The total number of *A. baumannii* GC1 and non-GC1 genomes that matched / not matched the unitigs in each dataset was used for calculation.

**Table S5. Rules obtained by the SCM models.** The table details the rule ID, the rule output from the SCM models, k-mer length and the location where the rule sequence matched in *A. baumannii* AYE genome (AN:CU459141.1) or *A. baumannii* ACICU genome (AN: CP000863.1) and the gene name corresponding to the genome region matched and the gene product. *A. baumannii* AYE and ACICU genomes were used as *A. baumannii* GC1 and non-GC1 references, respectively, to locate the k-mer sequences. The prefix “REGION” was used when the match occurred either in an intergenic region or in a combination of intergenic region and a gene.

**Table S6. Analysis of the SCM rules matches within the Dataset 1 and 2.** The table details the rule ID of the 100 larger k-mer sequences targeted by the SCM rules, the number of *A. baumannii* GC1 and non-GC1 genomes typified by MLST that matched the rule within the Datasets 1 and 2 using blastn and the p-values of Fisher’s exact test using a significance level of 0.05. Fisher’s exact test was calculated in R considering the nominal variables “Dataset Source” (Dataset 1 or Dataset 2) and “Matched” (Yes or No). The total number of *A. baumannii* GC1 and non-GC1 genomes that matched / not matched the rules in each dataset was used for calculation.

**Table S7. Prediction metrics on test dataset partitions from the Dataset 1 using the best performing SCM models.** Correlation between MLST typing and model prediction was calculated using Fisher’s exact test in R using a significance level of 0.05. The nominal variables “MLST Typing” and “Prediction” were considered during Fisher’s exact test calculation. The variable “MLST Typing” represented the genomes typed as *A. baumannii* GC1 (positive label) or *A. baumannii* non-GC1 (negative label) by MLST technique (true class). On the other hand, the variable “Prediction” represented the genomes predicted to be *A. baumannii* GC1 or non-GC1 by the SCM model (predicted class). We used the number of True Positives (TP), False Positives (FP), False Negatives (FN) and True Negatives (TN) in 2x2 contingency table. Null hypothesis used to evaluate the correlation between MLST typing and model prediction was: “True class (MLST Typing) and Predicted class are independent, knowing the value of one variable does not help to predict the value of the other variable”.

